# Shared genomic features of oral bacterial habitat specialization

**DOI:** 10.64898/2026.10.17.759843

**Authors:** Abish S. Stephen, Alan I. B. Vallavaraj

## Abstract

The oral microbiome exhibits spatial organization, with related bacteria occupying different surfaces despite continual dispersal between habitats. Whether this ecological partitioning is accompanied by genomic functions that recur across evolutionarily divergent lineages remains unclear. Here we compared plaque- and mucosa-associated bacteria representing 47 human microbial taxa, 13 genera and six phyla, using 1,121 reference genome assemblies and habitat-affinity estimates assembled from six oral microbiome abundance summaries. Across 13 genera, a genus-balanced screen of 13,917 nonredundant genomic profiles prioritised 14 exploratory annotations with nominal associations in the same direction before and after adjustment for genome composition with none passing genome-wide family-wise error correction. Proteins carrying recurrent domains belonged to multiple sequence clusters. Closely spaced, divergently oriented HxlR–SnoaL_2 gene pairs occurred in 66 genomes across four genera, although overall HxlR–SnoaL_2 linkage was not significantly associated with plaque preference. Using source-family-balanced habitat scores, nested cross-validation with entire genera held out supported prediction of habitat preference (macro-average within-genus AUC, 0.821; empirical P = 0.010). Predictive performance was lower after adjustment for genome size, GC content and coding-sequence number (AUC, 0.776; empirical P = 0.059). These findings reveal partial recurrence of habitat-associated genomic annotations amid lineage-dependent protein repertoires and genetic arrangements. They provide a comparative framework for distinguishing broadly shared ecological genomic signatures from alternative, potentially lineage-specific routes to oral habitat specialization.

## Introduction

The mouth contains a collection of microbial habitats in unusually close proximity. Non-shedding tooth surfaces support persistent, structured biofilms, whereas the tongue dorsum and buccal mucosa are epithelial environments with distinct surfaces, turnover dynamics, nutrient inputs and host interactions. These local differences generate reproducible ecological patterns: related bacterial taxa can be abundant at one oral site yet uncommon at another, even though saliva provides continual opportunities for dispersal [1–4]. Such biogeographic partitioning implies that closely related microorganisms can achieve different degrees of success in neighbouring environments, possibly due to specific biological traits associated with their distributions.

The site-specialist hypothesis has established an influential framework for understanding oral microbial biogeography [1]. Pangenomic analyses have demonstrated that species- and strain-level habitat preferences can be associated with differences in gene repertoires, including functions involved in intermediary metabolism, transport, cell-surface organization and stress responses. Studies of *Rothia* and *Haemophilus parainfluenzae* exposed cryptic habitat-associated populations within familiar oral taxa [5]; analyses of Saccharibacteria and other candidate phyla identified genetic correlates of tongue and plaque partitioning [6]; and work on Streptococcus, *Veillonella, Gemella, Haemophilus/Aggregatibacter* and Neisseriaceae further demonstrated that ecological differentiation extends across a range of oral lineages [7–11]. These investigations have been especially informative about variation within a genus, family or selected ecological group. However, their individual functional associations do not by themselves establish whether comparable genomic signals are recovered repeatedly across bacteria with widely separated evolutionary histories.

Different bacteria may exploit homologous biochemical systems, recruit different enzymes to a common reaction, alter regulation of ancestral pathways or acquire accessory functions through gene exchange. Equally, superficially similar Pfam or COG assignments may conceal divergent sequences and different substrates, whereas tightly linked genes may be inherited together without each contributing to the ecological phenotype. Distinguishing these possibilities requires a framework that considers habitat association across multiple genera, treats species rather than available genome assemblies as the ecological observations, and separates functional recurrence from common ancestry, genome composition and annotation redundancy. Our earlier oral ecological and biofilm studies likewise illustrate how changes in microbial community composition can be linked to molecular traits without identifying a uniquely causal function from association alone [12,13].

We therefore asked whether genomic signatures of plaque versus mucosal preference recur across unrelated oral bacterial genera, or whether similar ecological outcomes are accompanied principally by lineage-specific changes in genomic content. We assembled a panel of 47 human microbial taxa (HMTs) from 13 genera – containing species level microbial taxa that showed contrasting affinities to dental plaque biofilms or tongue and buccal mucosa. Species-level ecological affinities were linked to annotated prevalence profiles from 1,121 genomes, allowing us to compare equivalent plaque–mucosa contrasts within genera rather than simply pool taxa across a shared evolutionary tree. We first screened broad genomic annotation layers for recurrent associations, then resolved selected Pfam domains into full-length protein sequence clusters and inspected their chromosomal neighbourhoods. Finally, we tested whether feature patterns learned from 12 genera could predict ecological preference in a thirteenth genus withheld from discovery and modelling. This design permits a direct assessment of both the repeatability and the limits of cross-lineage genomic signatures, while leaving causal mechanisms as hypotheses for subsequent experimental investigation.

## Results

### Habitat specialization is repeatedly represented across divergent bacterial lineages

To examine genomic correlates of oral habitat preference across substantial evolutionary distance, we assembled 47 HMTs representing 13 genera and six bacterial phyla (Fig. 1a). Each genus contributed at least one plaque-associated and one mucosa-associated taxon, enabling comparisons within phylogenetically related groups while retaining a broad cross-lineage perspective. The final panel comprised 24 plaque-affiliated and 23 mucosa-affiliated taxa, supported by 1,121 reference assemblies. Between five and 50 assemblies were available per HMT (median, 19). Of the 47 taxa, 32 met the more stringent ecological-confidence definition and 15 met a moderate-confidence definition (Supplementary Table 1).

**Fig. 1.**
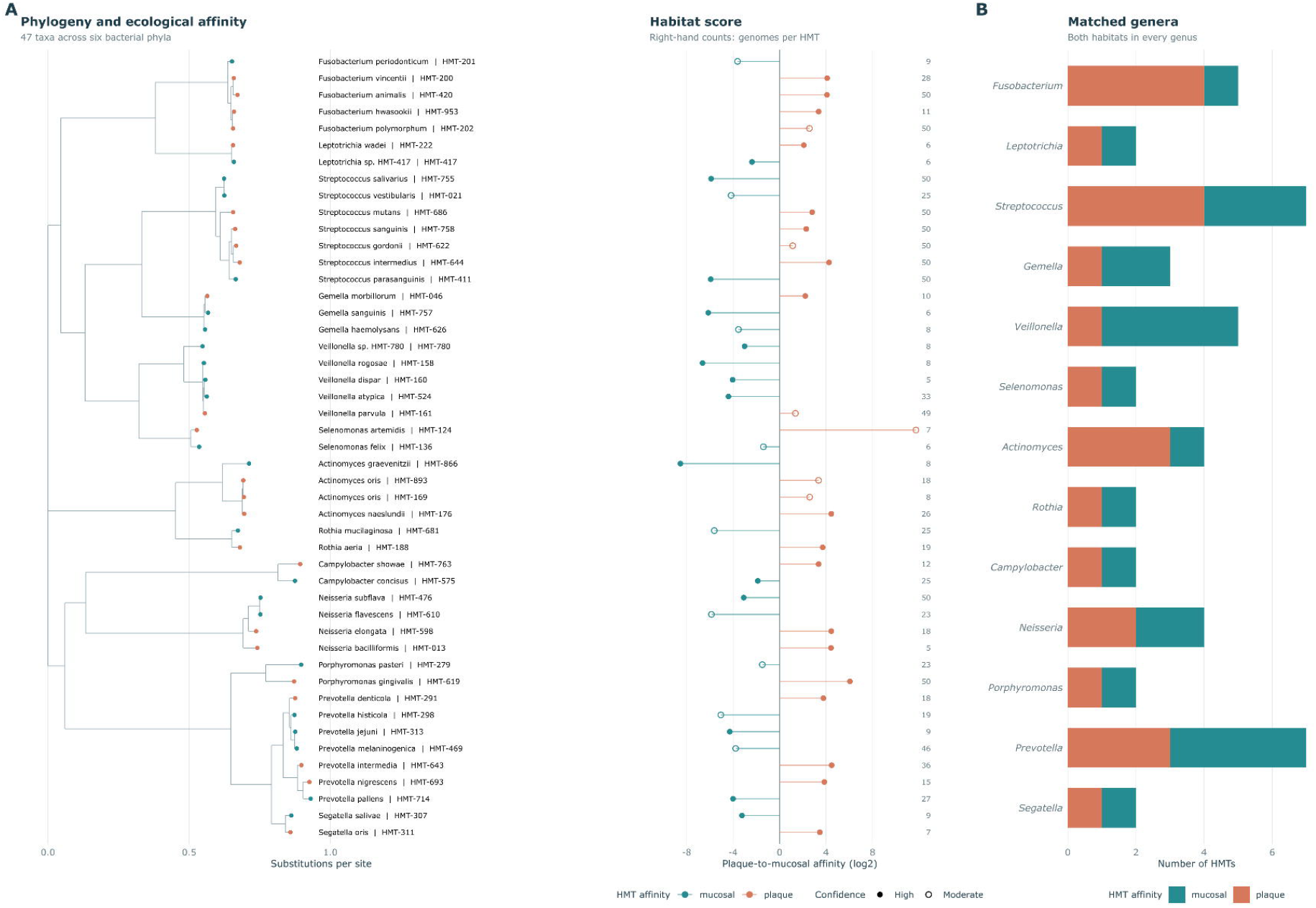
Oral habitat specialization is represented across 13 bacterial genera spanning six phyla. **a**, Branch-length phylogram inferred from 18 conserved marker proteins for the 47 selected Human Microbial Taxa (HMTs). Tip labels are accompanied by plaque-associated or mucosa-associated ecological classifications and the median source-derived plaque-to-mucosa log_2_ habitat-affinity score, calculated using subgingival plaque (SUBP) and supragingival plaque (SUPP) relative to buccal mucosa (BMU) and tongue dorsum (TDO). Solid symbols indicate high-confidence assignments (*n* = 32); open symbols indicate moderate-confidence assignments (*n* = 15). The right-hand track gives the number of reference genomes represented by each HMT (1,121 in total). **b**, Numbers of selected plaque- and mucosa-associated HMTs within each of the 13 genera; all genera contribute both ecological categories. The tree was display-rooted, and no ancestral habitat reconstruction or independently replicated evolutionary transition is inferred.

For each taxon, oral habitat preference was derived from six eHOMD-linked abundance summaries using the relative abundance in supragingival and subgingival plaque compared with tongue dorsum and buccal mucosa. The median source-specific log-ratio defined an ecological-affinity score, with positive values indicating greater relative plaque abundance and negative values indicating greater relative mucosal abundance (Methods). Because the underlying summaries partly reflect related sequence collections, they were treated as complementary ecological estimates rather than six independent biological cohorts. These scores describe relative site affinity and are not a measure competitive fitness, absolute occupancy, or habitat adaptation experimentally.

An 18-marker maximum-likelihood phylogeny separated the selected taxa into the expected major lineages while revealing contrasting habitat preferences within several relatively shallow branches (Fig. 1a). The genus-matched ecological design is therefore not driven by a single phylogenetic division between plaque and mucosal taxa. Rather, preference for these two broad habitat classes is distributed among organisms of substantially different ancestry. The adjacent ecological tracks in Fig. 1 illustrate this distinction more directly than a taxonomic comparison alone: taxa sharing genus membership may occupy opposite ends of the plaque–mucosa affinity gradient. This repeated representation provides the comparative opportunity for asking whether analogous genomic functions accompany similar ecological preferences in different genera.

The study was intentionally conditioned on strong habitat preferences and sufficient public genome representation (Fig. 1b). As a result, the 47 taxa should not be interpreted as a random sample of the oral microbiome or as a test of the prevalence of specialization among all oral bacteria. An additional limitation is the broad pooling of tongue and buccal mucosa, whose ecological pressures differ. We therefore retained the continuous affinity information and evaluated source-family and site-definition sensitivity analyses instead of equating all oral mucosal environments.

### A genus-balanced genomic atlas reveals recurrent but exploratory functional associations

We next asked whether genomically encoded functions showed a common direction of difference between plaque-associated and mucosa-associated taxa across the 13 genera. Genome-derived annotations were aggregated to HMT-level prevalence or annotation-derived scores, so that genomes contributed information about a taxon’s repertoire without being treated as independent ecological replicates. The complete representation contained 13,917 distinct 47-HMT profiles after collapsing profiles with identical distributions, comprising sequence clusters, Pfam domains, COG categories, enzymes, reactions and KEGG pathway/module annotations (Methods; Fig. 2a). For each profile, we calculated a plaque-minus-mucosa difference within each genus and assessed the coherence of direction across genera with 4,096 within-genus label permutations. The analysis was repeated after accounting for genome size, GC content and predicted coding-sequence number.

**Fig. 2.**
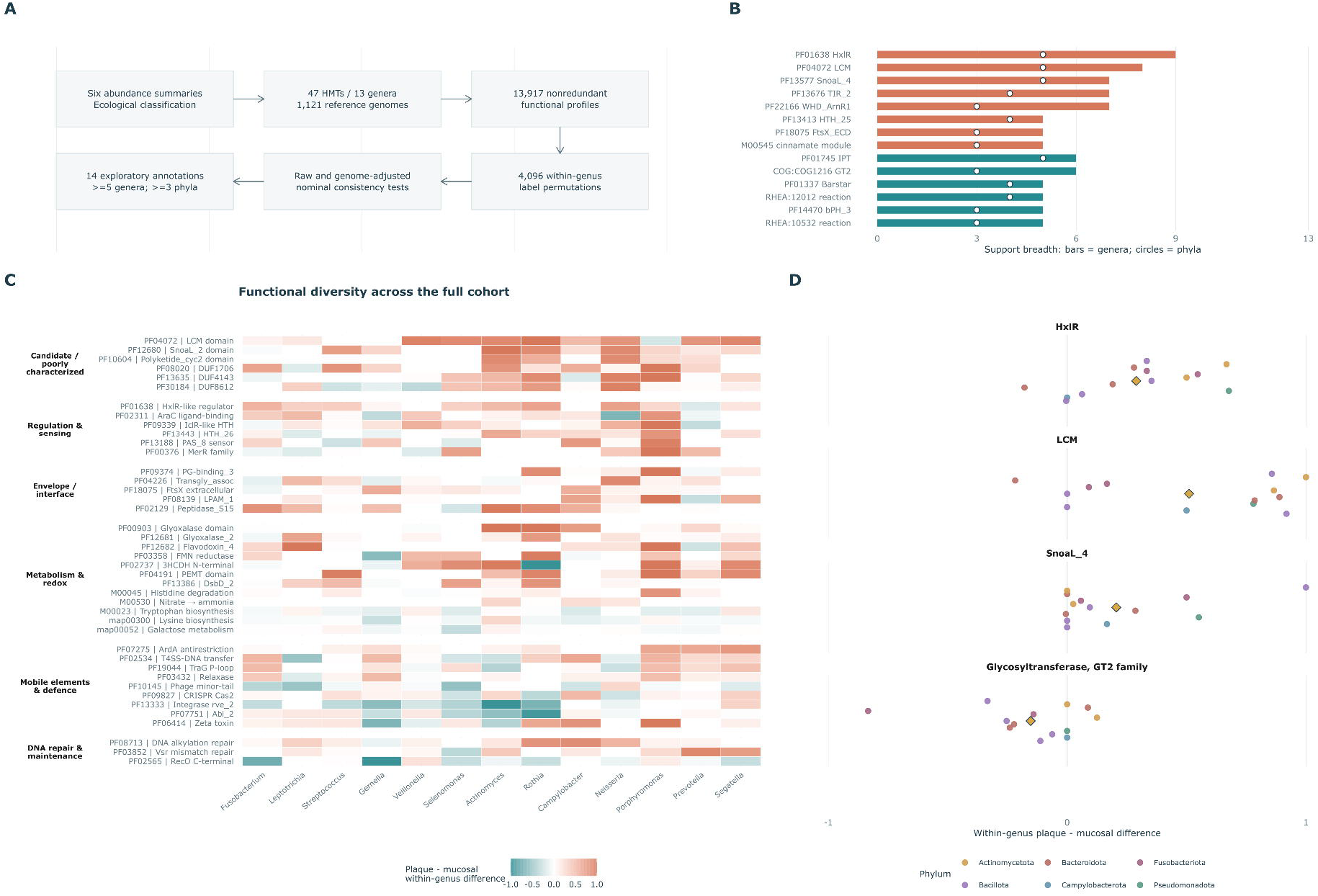
Genus-balanced recurrence and heterogeneity of genomic annotation profiles associated with oral habitat preference. **a**, Six source-specific oral abundance summaries informed ecological affinities for 47 HMTs, while 1,121 assemblies informed HMT-level genomic prevalence or annotation-derived scores. After exact-profile deduplication, 13,917 profiles were compared using equal-weight genus plaque-minus-mucosa contrasts and 4,096 within-genus label permutations, before and after genome size, GC content and CDS count adjustment. A total of 119 profiles reached nominal consistency *P* < 0.05 in the original screen, 23 were nominal in both screens, and 14 met the additional requirement for concordant direction and support from at least five genera and three phyla in each analysis. No individual profile survived all-feature maxT family-wise correction. **b**, Genus and phylum support for the 14 exploratory candidates. Bars show supporting genera and open circles show represented phyla, using the smaller count from the unadjusted and adjusted analyses at each taxonomic rank. Support requires a within-genus difference in the candidate’s direction exceeding the feature-specific effect threshold (Methods), rather than a separately significant genus-level test; the supporting taxa need not be identical between analyses. **c**, A separately curated 41-feature descriptive atlas showing heterogeneity of raw genus-level contrasts across six annotation themes; the 41 features were not selected by a formal relaxed *P*-value cutoff. **d**, Unadjusted plaque-minus-mucosa contrasts for four representative annotations (HxlR, LCM, SnoaL_4 and COG1216/GT2). Dots denote genera and diamonds equal-genus means. Binary domain prevalence, enzymatic-reaction labels and pathway/module annotation scores are distinct forms of evidence and do not confirm biochemical activities.

At a nominal threshold of *P* < 0.05 for the genus-level direction-consistency statistic, 119 profiles showed support in the unadjusted scan, of which 23 also showed nominal support after adjustment for genome composition. Requiring concordant overall direction between analyses, together with within-genus differences exceeding the specified effect-size threshold in at least five genera spanning at least three phyla in each analysis, yielded 14 exploratory candidate annotations (Fig. 2a,b; Supplementary Table 2). Eight were plaque-oriented and six were mucosa-oriented. No individual feature passed the all-profile maximum-statistic family-wise error correction in either analysis. These annotations therefore constitute a shortlist for further investigation, with nominal rather than genome-wide statistical support.

The plaque-oriented annotations included the HxlR-family domain PF01638, supported in nine genera across five phyla, and the LCM domain PF04072, supported in eight genera across five phyla. PF13577 (SnoaL_4) was supported in seven genera across five phyla, whereas PF13676 (TIR_2) and PF22166 (WHD_ArnR1) were each supported in seven genera, spanning four and three phyla, respectively. The remaining plaque-oriented annotations were PF18075 (FtsX extracellular domain), PF13413 (HTH_25) and KEGG module M00545 (trans-cinnamate degradation). The six mucosa-oriented annotations were PF01745 (IPT/MiaA), COG1216 (glycosyltransferase family 2), PF01337 (Barstar), PF14470 (bPH_3), RHEA:12012 (thiamine kinase reaction) and RHEA:10532 (tryptophan synthase reaction). Supporting counts reflect the direction and magnitude of within-genus differences, rather than separate statistically significant associations within each genus. The unadjusted and adjusted genus-level effects are provided in Supplementary Table 4.

The relationships were not uniform. A broader, separately curated 41-feature descriptive atlas displayed differences spanning regulatory proteins, poorly characterized domains, cell-envelope functions, metabolism, mobile genetic elements and DNA maintenance (Fig. 2c; Supplementary Table 3). Some genera differed across several of these annotation groups, whereas others showed relatively restricted or reversed patterns. This wider atlas was chosen after examining the data to illustrate the range of plausible functional changes and to help describe heterogeneity but with no additional discovery-level significance. Site-focused comparisons retained the expected direction for all 14 annotations across supragingival plaque–tongue, subgingival plaque–tongue and supragingival plaque– buccal contrasts, although nominal statistical support varied (Supplementary Table 14). These analyses are best interpreted as internal sensitivity checks rather than external replication.

Comparison with previous work places these associations in a broader ecological context. A recent study identified a leucine carboxyl methyltransferase annotation as a candidate marker of the plaque-associated ecotype of *Porphyromonas pasteri* [14], consistent with the plaque-oriented LCM signal in our analysis. Conversely, the mucosa-oriented COG1216 signal identifies an association at the level of a broad glycosyltransferase annotation, without resolving the substrates or cellular roles of the underlying proteins. Shared annotations can therefore reveal recurring ecological patterns while leaving their mechanistic basis unresolved. This distinction motivated our examination of the protein sequences and genomic contexts underlying the domain-level signals. An annotation-level evidence audit for all 14 candidates, distinguishing observed sequence evidence from untested mechanistic hypotheses, is provided in Supplementary Table 11.

### Similar domain annotations are embedded in different protein repertoires

To resolve the protein contexts underlying recurrent annotations, we linked the selected Pfam domains and related families to their full-length predicted proteins and MMseqs2 sequence clusters, defined at 50% sequence identity and 80% coverage (Fig. 3). A domain can occur in otherwise divergent proteins, while several Pfam models can annotate the same protein or overlap within it. We therefore asked how far the breadth of a domain across genera was retained at the level of full-length sequence clusters.

**Fig. 3.**
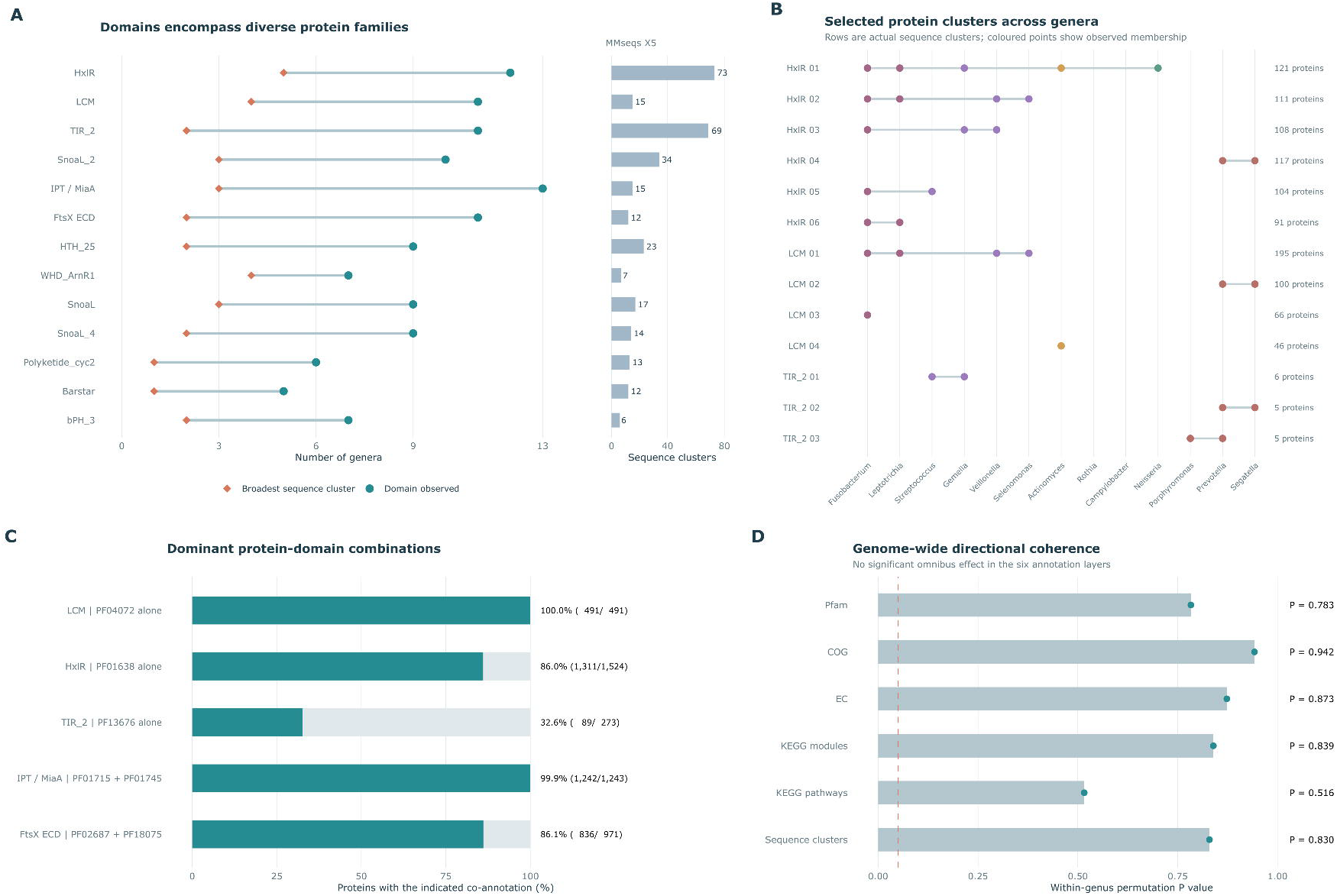
Recurrent **Pfam** domains **occur in heterogeneous protein** sequence groups. **a**, Genus breadth of selected domains and their broadest full-length sequence clusters, alongside numbers of sequence clusters. Clustering used 50% identity and 80% coverage. **b**, Genus membership of selected HxlR, LCM and TIR_2 clusters. **c**, Proportions of proteins carrying the indicated Pfam co-annotations. Co-annotation does not establish domain order, separate structural domains or common catalytic activity. **d**, One-sided permutation P values for repertoire-wide directional coherence across Pfam, COG, EC, KEGG-module, KEGG-pathway and sequence-cluster profiles. Tests used genome-adjusted profiles without feature-variance scaling and 5,000 within-genus label permutations. None showed significant coherence. Protein and assembly counts are descriptive; HMTs form the ecological observations.

The domain and protein views differed substantially. PF01638 (HxlR) occurred in 1,524 predicted proteins from 12 genera, distributed among 73 sequence clusters; the most widely distributed cluster spanned five genera. PF04072 (LCM) occurred in 491 proteins from 11 genera, divided among 15 clusters, with the broadest cluster spanning four genera. PF13676 (TIR_2) occurred in 273 proteins from 11 genera and 69 clusters, none spanning more than two genera (Fig. 3a,b; Supplementary Table 5). Thus, the recurrence of a domain across lineages often encompassed several distinct protein sequence groups. Pfam co-annotations further complicated the relationship between a domain label and a protein’s function. Some proteins carried several assignments, including models that overlapped within the same sequence region. For example, SnoaL_2 (PF12680) and SnoaL (PF07366) frequently annotated overlapping regions of the same protein. Their co-occurrence therefore cannot be treated as evidence for two separate genes or activities. The co-annotation patterns in Fig. 3c describe protein contexts; biochemical substrates and interaction partners remain to be established.

We next asked whether these selected associations formed part of a broader, coordinated shift in genomic content. After adjustment for genus, genome size, GC content and CDS count, we compared plaque-minus-mucosa effect vectors across six annotation layers: Pfam, COG, EC, KEGG modules, KEGG pathways and sequence clusters. None showed significant repertoire-wide directional coherence across the 13 genera, either without feature scaling (permutation P = 0.516–0.942; Fig. 3d) or after feature scaling (P = 0.747–0.937; Supplementary Table 12). These tests provided no evidence for a broadly aligned shift across the annotated repertoires, although they do not exclude a smaller set of recurring associations within otherwise heterogeneous genomic backgrounds. Together, these analyses separate recurrence of an annotation from recurrence of a particular protein. The same domain label can encompass different full-length sequences and co-annotation patterns across lineages - shared ancestry, gene exchange and differential gain or loss could all contribute to these distributions, but domain prevalence alone cannot distinguish their contributions.

### Conserved chromosomal orientation coexists with lineage-specific genomic arrangements

We then examined whether selected domain-bearing genes recurred in similar genomic arrangements. Among **the** candidate pairs examined within 3 kb on the same contig, HxlR (PF01638) and SnoaL_2 (PF12680) were linked in **67 genomes representing 11 HMTs** and **five genera**: *Actinomyces, Porphyromonas, Prevotella, Streptococcus* and *Neisseria* (Fig. 4a,b; Supplementary Table 6). In 66 genomes across the first four genera, the genes were divergently oriented and separated by 73–226 bp, with a median gap of approximately 140 bp. The single *Neisseria* example instead had a 2,124-bp gap and a same-strand orientation.

**Fig. 4.**
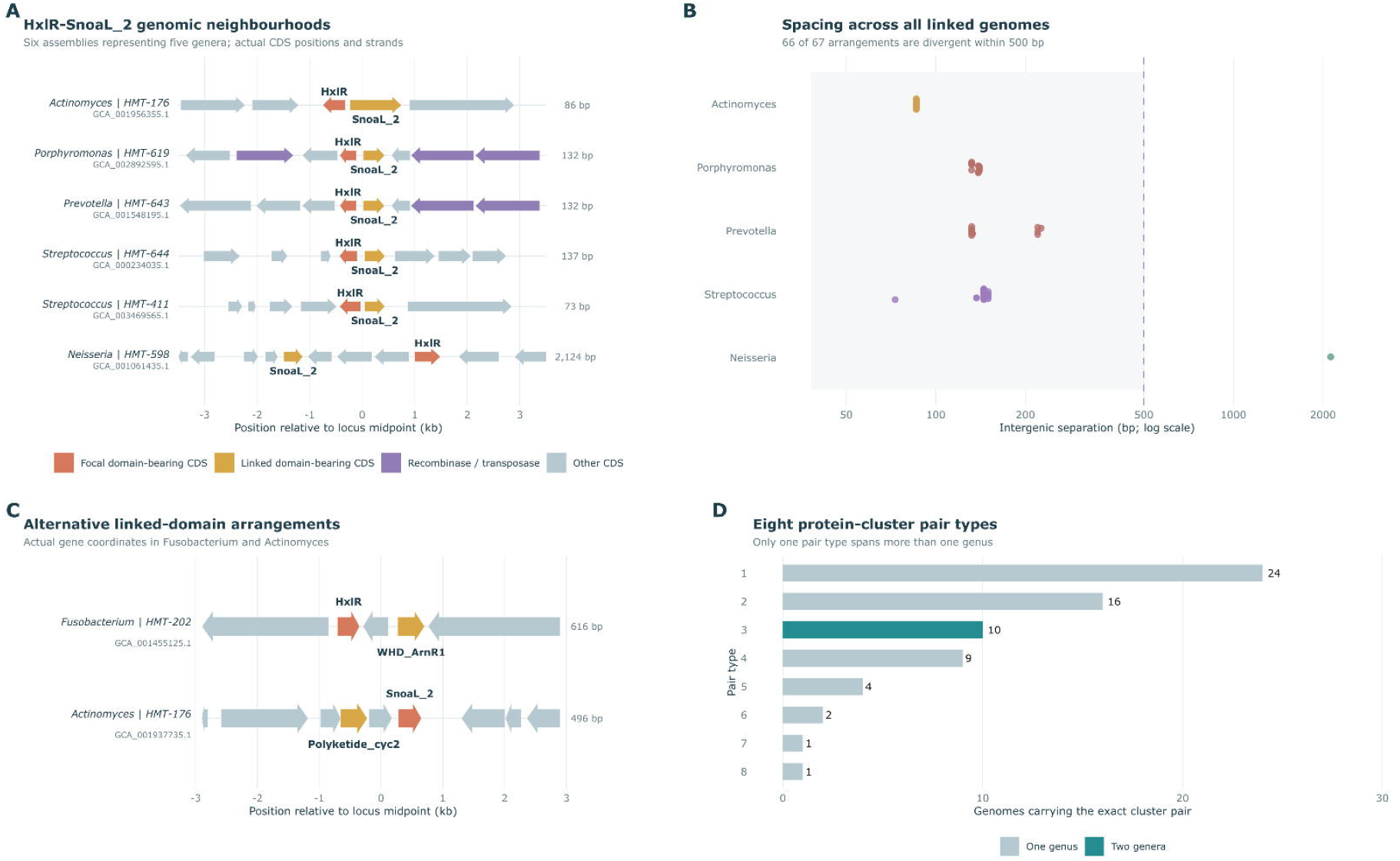
Recurrent gene orientation and variable sequence-cluster combinations among linked candidate domains. **a**, Prokka GFF-based gene-neighbourhood tracks for representative linked HxlR (PF01638) and SnoaL_2 (PF12680) loci. CDS extents and strand orientations are based on actual reference assembly coordinates; genome tracks are displayed relative to the central linked genes. b, Distribution of intergenic separations and orientations across the 67 HxlR–SnoaL_2 linked genomes from five genera. In 66 genomes across four genera, the two genes were divergent and separated by at most 500 bp; the isolated *Neisseria* example was same-strand with a 2,124-bp gap. c, Representative local architectures involving HxlR–WHD_ArnR1 (38 genomes, *Fusobacterium*) and SnoaL_2–Polyketide_cyc2 (35 genomes, *Actinomyces*). d, Eight full-length sequence-cluster pair types occurred among the 67 HxlR–SnoaL_2-linked genomes; only one pair type extended across more than one genus. The two-sided genus-balanced linkage test used HMT-level prevalence across all 47 HMTs, with linkage defined by the full 3-kb criterion (*P* ≈ 0.217; Supplementary Table 6b); it was not a separate test of the 66 divergent arrangements. Recurrence of gene orientation does not establish shared regulation or an evolutionary mechanism.

In the closely spaced pairs, the predicted transcriptional directions point away from a short intergenic region. This arrangement is compatible **with** local regulatory coupling, although promoter activity and shared regulation have not been demonstrated. Tracks drawn from the GFF coordinates showed that divergent orientation persisted despite variation in the surrounding genes (Fig. 4a). Linkage also occurred in mucosa-associated taxa. A genus-balanced test of HxlR–SnoaL_2 carriage, using HMT-level linkage prevalence across all 47 HMTs and the broader 3-kb linkage definition, did not detect a significant association with plaque preference (two-sided permutation *P* ≈ 0.217; Supplementary Table 6b). The recurrent architecture and its ecological association are therefore distinct observations.

The linked proteins were also heterogeneous. The 67 genomes contained **eight combinations of HxlRbearing and SnoaL_2-bearing** sequence **clusters**, only one of which occurred in more than one genus (Fig. 4d). Domain adjacency was therefore more broadly shared than any individual pair of full-length sequence clusters. This pattern could arise through inheritance of an ancestral arrangement, replacement of its components or repeated assembly of related genes. The arrangement alone does not establish which evolutionary route occurred. Other examined domain pairs were restricted to individual genera within the study panel. HxlR and WHD_ArnR1 occurred within 3 kb in 38 Fusobacterium genomes, while SnoaL_2 and PF10604 (Polyketide_cyc2) were linked in 35 *Actinomyces genomes* representing three HMTs (Fig. 4c). These arrangements illustrate how similar domain annotations can occur in different local gene contexts. The Polyketide_cyc2 label does not establish a polyketide pathway, and adjacency alone does not demonstrate co-transcription. Genomic context therefore provides hypotheses about the systems in which candidate proteins operate. Recurrent spacing and orientation may reflect regulatory constraints or joint inheritance, while linkage may cause a domain to track another gene that contributes to habitat preference. These possibilities remain to be distinguished by evolutionary and functional analyses.

### Close-relative comparisons reveal consistent differences and informative reversals

To examine the same ecological contrasts among close relatives, we identified eight terminal sister pairs with opposite habitat affinities in the 18-marker tree (Extended Data Fig. 1; Supplementary Table 8). The pairs represented *Leptotrichia, Gemella, Selenomonas, Rothia, Campylobacter, Porphyromonas, Prevotella* and *Segatella*. Seven sister nodes had 100% bootstrap support; the *Gemella* node had 59% and provided a less secure phylogenetic comparison. Because all pairs came from the primary 47-HMT panel, this analysis assesses sensitivity to relatedness rather than independent replication. LCM (PF04072) was more prevalent in the plaque-associated taxon in six pairs, unchanged in one and more prevalent in the mucosa-associated *Porphyromonas* taxon. SnoaL_2 (PF12680) was plaque-oriented in six pairs and unchanged in two. These comparisons recover recurring directions while retaining informative exceptions. In particular, the *Porphyromonas* LCM contrast differs from the plaque-ecotype association reported within *P. pasteri* [14]. The two comparisons involve different taxonomic resolutions and cannot establish a single ecological direction for the entire domain family.

Broader functional categories showed similar dependence on the comparison and representation. After adjustment for genus and genome covariates, scores for the primary COG2024 cell wall, membrane and envelope category M were higher on the mucosal side in all eight sister pairs (exact two-sided sign-flip *P* = 0.0078; category-wise maxT *P* = 0.109). Scores for translation-associated category J were higher on the plaque side in seven pairs (*P* = 0.0156; maxT *P* = 0.094). Neither category passed family-wise correction (Supplementary Table 9). These exploratory category-level patterns complement the individual-feature contrasts without establishing a shared functional programme across lineages. The sister-pair analysis thus places the cross-genus patterns in a more local phylogenetic context. It reveals both agreement and reversals within the same study panel, while the sampled sister relationships do not establish independently repeated evolutionary transitions. Extended Data Fig. 1 shows the domain-level sister-pair contrasts; the COG category tests are reported in Supplementary Table 9.

### Functional repertoires predict habitat preference in genera excluded from training

We finally asked whether genomic profiles could predict habitat affinity in a genus excluded from training. Each of 13 outer cross-validation folds withheld one entire genus. Within the remaining 12 genera, feature ranking, feature-set size and ridge-regularization strength were selected through inner leave-one-genus-out validation (Fig. 5a; Methods). The held-out genus did not contribute to these supervised selection and fitting steps. Predictions were made for HMTs, and performance was measured as the mean within-genus AUC: the proportion of plaque–mucosa taxon pairs ranked in the expected order, averaged equally across genera. Using the source-family-balanced habitat-affinity score, the model achieved a macro within-genus AUC of 0.821 (95% genus-bootstrap interval, 0.641– 0.962; Fig. 5b–d; Supplementary Tables 7 and 10). A null distribution generated by 200 within-genus phenotype permutations, each repeating the nested workflow, gave an empirical *P* = 0.00995. The original median-based outcome gave a similar AUC of 0.808 (95% interval, 0.615–0.962; *P* = 0.00995). Performance nevertheless varied across genera. For the source-family-balanced model, *Selenomonas* showed reversed ordering (AUC 0), *Fusobacterium* and *Gemella* had AUCs of 0.5, and *Prevotella* had an AUC of 0.667. The overall average therefore describes transfer across the panel rather than uniform performance in every genus.

**Fig. 5.**
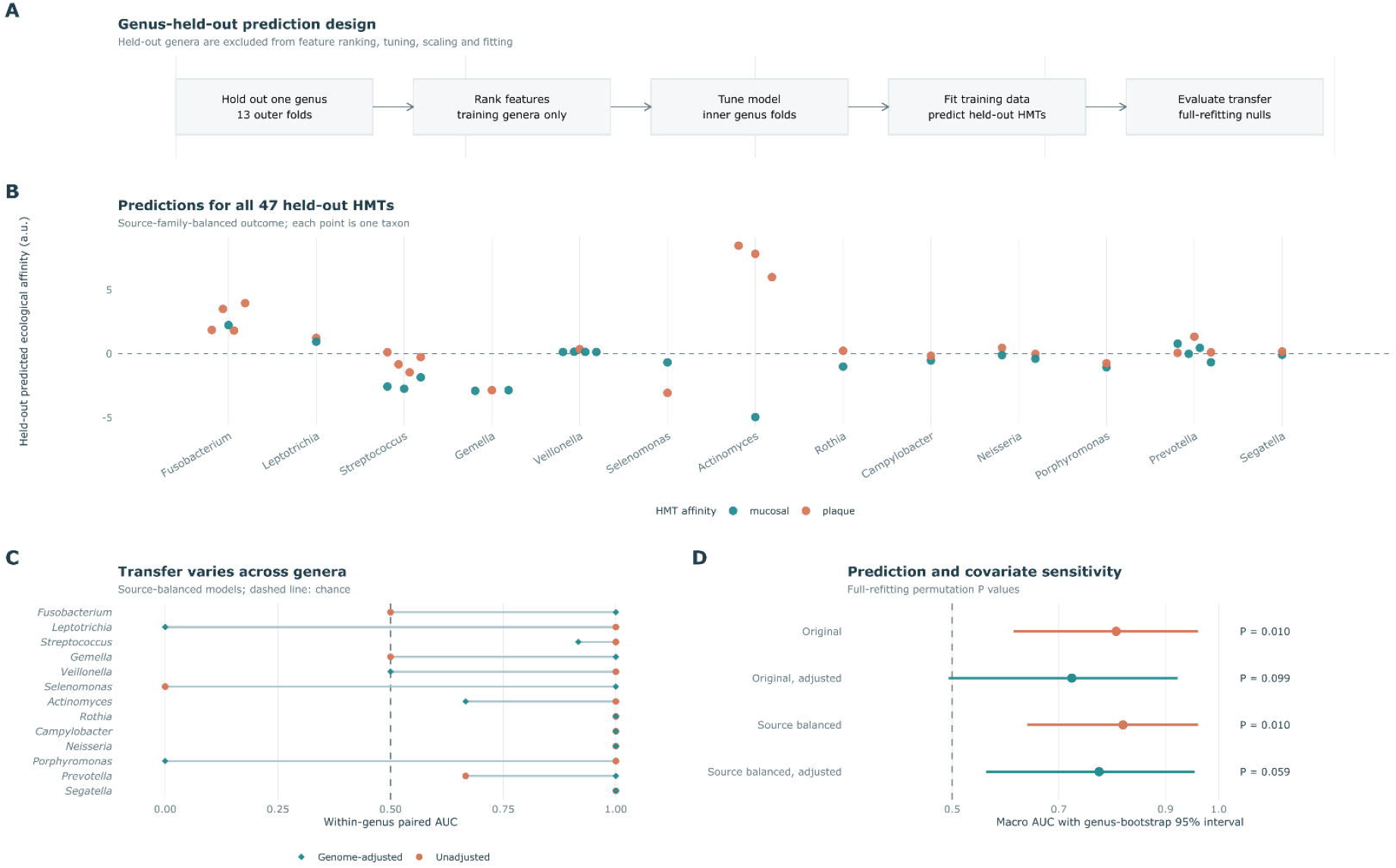
Genus-held-out prediction reveals transferable genomic information with variable performance and genome-content sensitivity. **a**, Fully nested leave-one-genus-out design: the excluded genus was not used for feature ranking, inner tuning or ridge-model fitting. **b**, Unadjusted source-family-balanced habitat-affinity predictions for all 47 HMTs, labelled by ecological class and grouped by held-out genus. Points are HMT-level values; reference genomes are not independent prediction samples. **c**, Genus-specific within-genus AUC for source-family-balanced predictions before and after genome-covariate residualisation. The dashed reference indicates chance-level rank discrimination (0.5). **d**, Macro within-genus AUC and 95% genus-bootstrap percentile intervals for original and source-family-balanced outcomes with or without adjustment for genome size, GC content and CDS count. The unadjusted genus-balanced models achieved AUC 0.808 and 0.821, respectively (*P* = 0.00995, 200 full-refitting within-genus permutations each). The corresponding adjusted models achieved AUC 0.724 (*P* = 0.0990) and 0.776 (*P* = 0.0594) under 100 full-refitting permutations each. AUC concerns within-genus ranking of selected specialists rather than prediction of absolute habitat abundance in an independent clinical sample.

To assess sensitivity to broad genome characteristics, we repeated the analysis after removing feature variation associated with genome size, GC content and CDS count, fitting this adjustment within each inner and outer training partition. The source-family-balanced model then achieved an AUC of 0.776 (95% genus-bootstrap interval, 0.564–0.955; empirical *P* = 0.0594 from 100 full-refit permutations). For the original outcome, the corresponding AUC was 0.724 (95% interval, 0.494–0.923; *P* = 0.0990). These lower point estimates and weaker permutation evidence leave the extent of prediction independent of broad genome properties unresolved (Fig. 5d; Supplementary Table 10). We also tested how predictions changed when selected annotation groups were excluded and the full nested workflow was retrained. For the original outcome, removing LCM changed AUC from 0.808 to 0.705; removing HxlR left it at 0.808; and removing four SnoaL-related annotations increased it to 0.833. Excluding LCM, HxlR and those four annotations together reduced AUC to 0.603. Excluding the 14 shortlisted annotations plus SnoaL (PF07366), SnoaL_2 (PF12680) and Polyketide_cyc2 (PF10604) reduced it to 0.577 (Supplementary Table 13). These post hoc exclusions describe the sensitivity of the modelling workflow to particular annotation groups. Correlated features and model retuning prevent the AUC changes from being assigned as independent or causal contributions of individual proteins.

Collectively, the prediction results show that genomic profiles contain habitat information that transfers across some genus boundaries in the selected panel. Transfer was uneven, and its independence from genome composition remains uncertain. This supports further investigation of combinations of genomic features while retaining the exploratory status of individual annotations.

## Discussion

Oral bacteria occupy neighbouring surfaces with different attachment opportunities, nutrient supplies and host interactions despite continual dispersal through saliva. Previous work has resolved habitat preferences and associated genomic differences within individual genera [1,5–11]. Here, we compared the same plaque–mucosa contrast across 13 genera and six phyla. The genus-balanced screen identified 14 nominally supported annotations, but none passed genome-wide family-wise correction. Collective genomic profiles nevertheless predicted relative habitat preference in held-out genera, with variable performance and weaker evidence after genome adjustment. The study therefore identifies both recurring genomic patterns and limits to their transfer across lineages. A shared ecological challenge may not have the same genetic solution in every lineage. Persistence on teeth or mucosal surfaces could involve related biological processes carried out by different proteins or regulated in different ways. The heterogeneous protein clusters and genomic contexts observed here are compatible with that possibility. However, the repertoire-wide tests did not establish a coordinated genomic shift, and the individual associations could also reflect ancestry, linked genes or genome composition. Distinguishing these explanations requires finer functional resolution than broad annotation labels provide.

Previous studies provide useful points of comparison. Metapangenomic analyses have linked population structure and gene repertoires to oral habitats in *Haemophilus parainfluenzae* and *Rothia* [5], with subsequent work extending site-specific genomic comparisons to other genera [8–11]. A leucine carboxyl methyltransferase annotation has been identified as a candidate marker of the plaque-associated *P. pasteri* ecotype [14]. Our plaque-oriented LCM signal extends this comparison across genera, while the reversed *Porphyromonas* sister-pair contrast shows why results at different taxonomic resolutions need not agree. The contribution of the present study is to assess the breadth and heterogeneity of such associations, rather than to assign a universal ecological role to a protein family. The HxlR–SnoaL_2 neighbourhood illustrates how a recurring arrangement can coexist with protein diversity. Closely spaced, divergently oriented genes occurred across four genera, but the linked proteins belonged to several sequence-cluster combinations. The short intergenic region is a plausible site of regulatory interaction, although neither shared promoter control nor a ligand response has been demonstrated in these oral homologues. Comparative motif and sequence analyses can guide experiments, but promoter reporters, DNA-binding measurements and ligand-response assays are needed to test whether these genes form a regulatory unit. The same caution also applies to the evolutionary interpretation of linkage. HxlR–WHD_ArnR1 and SnoaL_2–Polyketide_cyc2 were restricted to *Fusobacterium* and *Actinomyces*, respectively, within the 13-genus panel and their distributions could reflect lineage-specific inheritance, acquisition or loss. The broader recurrence of divergent HxlR–SnoaL_2 pairs could likewise arise through inheritance or gene exchange, with local regulatory constraints preserving orientation. Comparisons of gene and species trees, flanking mobile elements and sequence-level boundaries are needed to separate these possibilities. Recurrent orientation alone cannot establish convergent evolution.

The unadjusted nested models ranked habitat affinities in held-out genera better than the within-genus permutation null, indicating that useful information was shared across some lineages. Performance was uneven: the source-family-balanced model reversed the *Selenomonas* contrast and gave chance-level ordering in *Fusobacterium* and *Gemella*. Genome adjustment produced lower AUC estimates and weaker permutation evidence, but this does not quantify the causal contribution of genome composition or establish a significant difference between models. Larger panels and independent ecological data, together with phylogenetic and genome-covariate baselines, would help determine how much predictive information is specific to the annotated functions.

Several features of the design limit generalization: The panel was selected for consistent habitat preferences and adequate genome representation, so it cannot establish how common these patterns are among oral generalists or poorly sampled taxa. The six abundance summaries share underlying data sources; balancing source families reduces unequal representation without making the summaries independent. Uneven genome sampling, non-oral isolation sources and incomplete metadata may influence estimated annotation prevalence. The 13 genera define the blocks of comparison rather than the 1,121 assemblies, and these genera also share evolutionary history. Most contain few sampled HMTs, limiting the resolution of within-genus prediction. Finally, broad and overlapping annotation labels constrain functional interpretation, and no individual profile passed genome-wide family-wise correction.

This study provides a framework for comparing oral habitat associations across bacterial lineages while separating annotation recurrence, protein identity and genomic organisation. It reveals candidate patterns that can guide functional work and collective genomic information that transfers unevenly across genera. Establishing which patterns reflect shared ecological mechanisms will require independent sampling and direct tests of the implicated proteins and regulatory systems.

## Methods

### Study design and taxon selection

The study was a retrospective comparative-genomics analysis of publicly available genomes and previously published oral abundance summaries. The primary panel comprised 47 HMTs assigned to 13 genera, each represented by both plaque-associated and mucosa-associated taxa. Taxa were selected for sufficient reference-genome availability and consistent ecological direction across the relevant source summaries. The panel included six phyla and 24 plaque-associated versus 23 mucosa-associated HMTs. The HMT was the biological observational unit for ecological comparisons. Genomes nested within an HMT were used to estimate genomic content and its prevalence, rather than counted as independent biological replicates.

### Construction of the ecological habitat-affinity phenotype

We obtained HMT-level abundance estimates from six eHOMD-linked source summaries corresponding to the Dewhirst, Eren V1–V3, Eren V3–V5, HMP MetaPhlAn and HMP RefSeq V1–V3 and V3–V5 resources [2,4,15]. Four oral environments were used: subgingival plaque (SUBP), supragingival plaque (SUPP), buccal mucosa (BMU) and tongue dorsum (TDO). For HMT *h* and source *s*, we defined

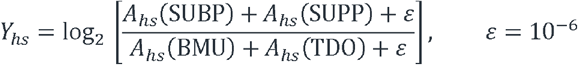

Here, *A* denotes the reported relative-abundance summary for the taxon and site. The original HMT phenotype, *Y*⍰, was the median of eligible source-specific scores, with positive values indicating plaque affinity and negative values indicating affinity for the combined tongue and buccal mucosal sites. Source records with identical non-zero four-site abundance profiles assigned to multiple HMTs within the same genus were excluded as potentially ambiguous. The HMP MetaPhlAn estimate for *Campylobacter* concisus HMT-575 was also excluded because of a source-marker inconsistency. High-confidence status required at least five contributing sources, at least four abundance-qualified sources, at least 80% directional agreement and |Y⍰| ≥ 1.5; moderate confidence required at least four sources, two abundance-qualified sources, at least 80% agreement and |Y⍰| ≥ 1.0. A source was abundance-qualified when the sum across the four focal sites was at least 0.1 on the supplied abundance scale. The resulting panel contained 32 high-confidence and 15 moderate-confidence HMTs.

To reduce unequal representation of related abundance summaries, we also calculated a source-family-balanced phenotype. For each HMT, source-specific scores were first summarised by their median within each of the Dewhirst, Eren and HMP families; the median of the available family medians defined the balanced score. Nested predictions were evaluated using both this score and the original median-based outcome. Additional sensitivity analyses used SUBP–TDO, SUPP–TDO and SUPP–BMU contrasts. Because these contrasts reuse the underlying abundance summaries, they were treated as internal sensitivity checks.

### Genome retrieval, quality control and taxonomic assignment

Publicly available genomic assemblies assigned to the selected HMTs were curated into a nonredundant genome manifest and quality assessed using the established workflow. The manifest was filtered for CheckM2-estimated completeness of at least 95% and contamination of no more than 5%, unique assembly accessions, and a maximum of 50 retained genomes per HMT [16]. The final dataset contained 1,121 genomes, with five to 50 genomes per HMT and a median of 19. Assembly provenance was retained at accession level, including metadata on the reported sample source and whether records were designated metagenome assembled. Of the retained genomes, 154 were flagged as MAG-derived; some others originated from extraoral samples or lacked a fully interpretable isolation-source description. These entries were not relabelled as oral isolates merely because their taxonomy corresponded to an oral HMT. Unequal genome representation and genome-source uncertainty are limitations of genomic prevalence estimation.

### Genome annotation and HMT-level genomic profiles

Genome annotations combined archived eHOMD Prokka outputs for 543 assemblies, documented as version 1.14.6, with annotations generated locally using Prokka version 1.15.6 for 578 assemblies [17]. COG and Enzyme Commission (EC) annotations were extracted from the Prokka tables. Predicted proteins were searched against Pfam-A using HMMER version 3.4 with family-specific gathering thresholds [18], and clustered using MMseqs2 easy-linclust with a minimum sequence identity of 50% and coverage threshold of 80% in coverage mode 0 [21]. Rhea reaction annotations were inferred from complete EC numbers using the archived EC-to-Rhea mapping, and KEGG pathway and module profiles were derived from EC-to-KEGG mappings downloaded on 30 September 2026 [19,20].

For binary annotations, HMT-level prevalence was calculated as the fraction of genomes within each HMT carrying the annotation. For KEGG pathways and modules, each genome received a score equal to the fraction of linked EC annotations detected, using only linked ECs represented in the study dataset as the denominator. These scores were averaged across genomes within each HMT; pathways and modules required at least two mapped ECs in the dataset. Accordingly, M00545 represented an annotation-derived EC-coverage score, rather than evidence of a complete trans-cinnamate degradation pathway or measured metabolic activity. The cross-genus screen retained profiles that varied across HMTs and were represented in at least two genera. Collapsing identical numerical profiles across the 47 HMTs reduced 17,694 eligible annotation profiles to 13,917 distinct profiles. Their representative labels comprised 5,745 sequence clusters, 4,850 Pfam families, 1,575 COGs, 1,124 Rhea reactions, 251 KEGG modules, 246 EC annotations and 126 KEGG pathways. Deduplication prevented identical distributions from being tested repeatedly, but do not make the retained profiles biologically independent. The feature dictionary and equivalence mappings preserve the links between representative profiles and their associated annotation labels.

### Phylogenetic reconstruction and sister-pair definition

Representative proteomes for the 47 HMTs were used to construct a species-level phylogeny based on 18 conserved translation-associated marker proteins. Markers were aligned using MAFFT [22], concatenated, and analysed by maximum likelihood with RAxML-NG [23]. Branch support was assessed with 200 bootstrap replicates. Terminal sister pairs were identified from the resulting tree, and pairs were retained for the phylogenetic sensitivity analysis when the two taxa differed in the sign of their ecological habitat-affinity score. Eight such opposing-habitat terminal pairs were recovered. Seven had 100% bootstrap support; the *Gemella* pair had 59%. The pairs were used as a sensitivity analysis within the **same** 47-HMT panel; terminal sister relationships are not implied as independent evolutionary transitions. For COG category sensitivity tests, each COG was assigned to its first COG2024 category, and category scores were the mean prevalence across mapped COG profiles. Twenty categories with at least ten mapped COGs and non-negligible raw and adjusted variance were retained.

Scores were adjusted for genus, genome size, GC content and CDS count; all 256 sign patterns across the eight sister-pair differences were enumerated, with maximum-statistic correction across the 20 categories (Supplementary Table 9).

### Genus-balanced feature contrasts and permutation testing

For genomic feature f and genus g, the fundamental effect was the difference in average HMT-level prevalence or score between taxa assigned to plaque and mucosal preference:

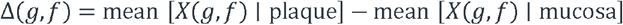

Each genus contributed one effect vector, irrespective of the number of HMTs or assemblies it contained. The direction-consistency statistic was the absolute difference between the numbers of genera with positive and negative effects exceeding an effect-size threshold. For each feature, this threshold was the larger of 0.05 and one-quarter of its standard deviation across HMTs in the unadjusted matrix, and the same threshold was used for adjusted contrasts. A studentized equal-genus mean-effect statistic was also calculated. Null distributions used 4,096 within-genus permutations of habitat labels, preserving group sizes. Adjusted profiles were residuals from regression on an intercept, genus indicators and HMT-level genome size, GC content and CDS count. For these screen tests, residualisation was performed once before label permutation.

Empirical *P* values included the observed configuration through a plus-one correction. For family-wise control, each permutation contributed the maximum statistic across all 13,917 profiles. No feature had an all-profile maxT *P* < 0.05. The exploratory shortlist required concordant signs of the mean genus effect, nominal direction-consistency *P* < 0.05 in both analyses, and at least five supporting genera spanning at least three phyla in each analysis. Reported genus and phylum counts were the smaller of the corresponding unadjusted and adjusted counts, calculated separately; the identities of supporting taxa could differ between analyses. The 41-feature descriptive atlas was curated after inspection of the data and was not defined by a relaxed significance threshold.

### Repertoire-wide tests of directional coherence

We compared plaque-minus-mucosa effect vectors across 13 genera using retained profiles from six annotation groups: Pfam, COG, EC, KEGG modules, KEGG pathways and sequence clusters. Profiles were analysed without scaling and after division by their across-HMT standard deviation, with a denominator floor of 0.1. Both representations were residualised against genus, genome size, GC content and CDS count. An omnibus statistic measured alignment through between-genus inner products, normalised by total squared effect magnitude and the number of other genera. One-sided empirical P values used 5,000 within-genus label permutations; Benjamini–Hochberg q values were calculated across the 12 group–scaling combinations. Results for all 12 representations are provided in Supplementary Table 12.

### Protein sequence clusters and genomic neighbourhood analysis

Pfam-bearing proteins were linked to their full-length predicted sequences, assembly accessions and HMT identities. MMseqs2 easy-linclust used minimum sequence identity 0.50, coverage 0.80 and coverage mode 0 [21]. We summarised cluster membership and taxonomic breadth and inspected co-annotation coordinates for overlapping Pfam assignments. Prokka GFF coordinates identified the nearest pair of distinct domain-bearing coding sequences on the same contig in each genome. Pairs separated by ≤3 kb were classified as linked, with overlapping coding sequences assigned zero separation. Gene order and strand defined divergent, convergent or same-strand orientation. HxlR– SnoaL_2 linkage prevalence within each HMT was compared between habitats using an equal-genus mean difference and 50,000 within-genus label permutations. This exploratory, two-sided test included all 47 HMTs and the full 67-genome linkage definition, without genome-wide correction. Accession-level linkage records and all HMT-level prevalence denominators are provided in Supplementary Tables 6 and 6b.

### Nested leave-one-genus-out prediction

Outer cross-validation withheld one genus at a time. Within training partitions, features were ranked by mean habitat-associated difference, between-genus dispersion and directional support above an effect threshold of 0.05. Features supported in fewer than two genera or showing negligible variation received zero ranking scores. Inner leave-one-genus-out validation selected among 5, 15, 40 or 100 features and ridge penalties of 10, 100 or 1,000. Selection maximised macro within-genus AUC, followed by Spearman correlation, lower mean absolute error, fewer features and lower penalty as tie-breakers. Ranking, standardisation and fitting were repeated within training folds; initial feature filtering and deduplication used the complete panel. Within-genus AUC measured the fraction of plaque–mucosa HMT pairs ranked correctly, with ties scored as 0.5; macro AUC averaged these values equally across genera. Both continuous outcomes used the original habitat labels. Null distributions jointly permuted outcomes and labels within genus and repeated the complete nested workflow: 200 permutations per unadjusted outcome and 100 per genome-adjusted outcome. Genome adjustment regressed features against genome size, GC content and CDS count separately within each training partition. Empirical P values used the plus-one correction. Ninety-five per cent percentile intervals used 50,000 bootstrap resamples of the 13 fixed genus-level AUCs, without model refitting.

### Post hoc feature exclusion and additional ecological sensitivity

We repeated nested prediction after excluding LCM, HxlR, four SnoaL-related annotations (PF07366, PF12680, PF13577 and PF10604), their specified combinations, or the 14 candidates plus PF07366, PF12680 and PF10604. These post hoc comparisons assess model sensitivity rather than independent or causal contributions (Supplementary Table 13). Site sensitivity used source-family-balanced scores for supragingival plaque–tongue, subgingival plaque–tongue, supragingival plaque–buccal mucosa, subgingival–supragingival plaque and tongue–buccal mucosa. Genus-centred Pearson correlations were tested using 10,000 within-genus permutations, with two-sided maximum-statistic correction across the 14 candidates within each contrast. The first three contrasts formed the principal plaque– mucosa summary (Supplementary Table 14). These analyses reuse the original abundance sources and provide internal sensitivity checks.

### Computing and reproducibility

Computational analyses used the Apocrita high-performance computing service at Queen Mary University of London. Statistical analyses were implemented in Python, and the five main figures and supplementary genomic neighbourhood atlas were generated in R. The accompanying computational package contains analysis scripts, fixed input datasets, reference outputs and software dependency specifications for the 13-genus study. Reproduction from these inputs verified the primary feature screen and observed nested prediction results against the archived outputs. Workflow documentation specifies execution instructions, external data requirements and the scope of these checks.

## Figure legends

**Extended Data Fig. 1.**
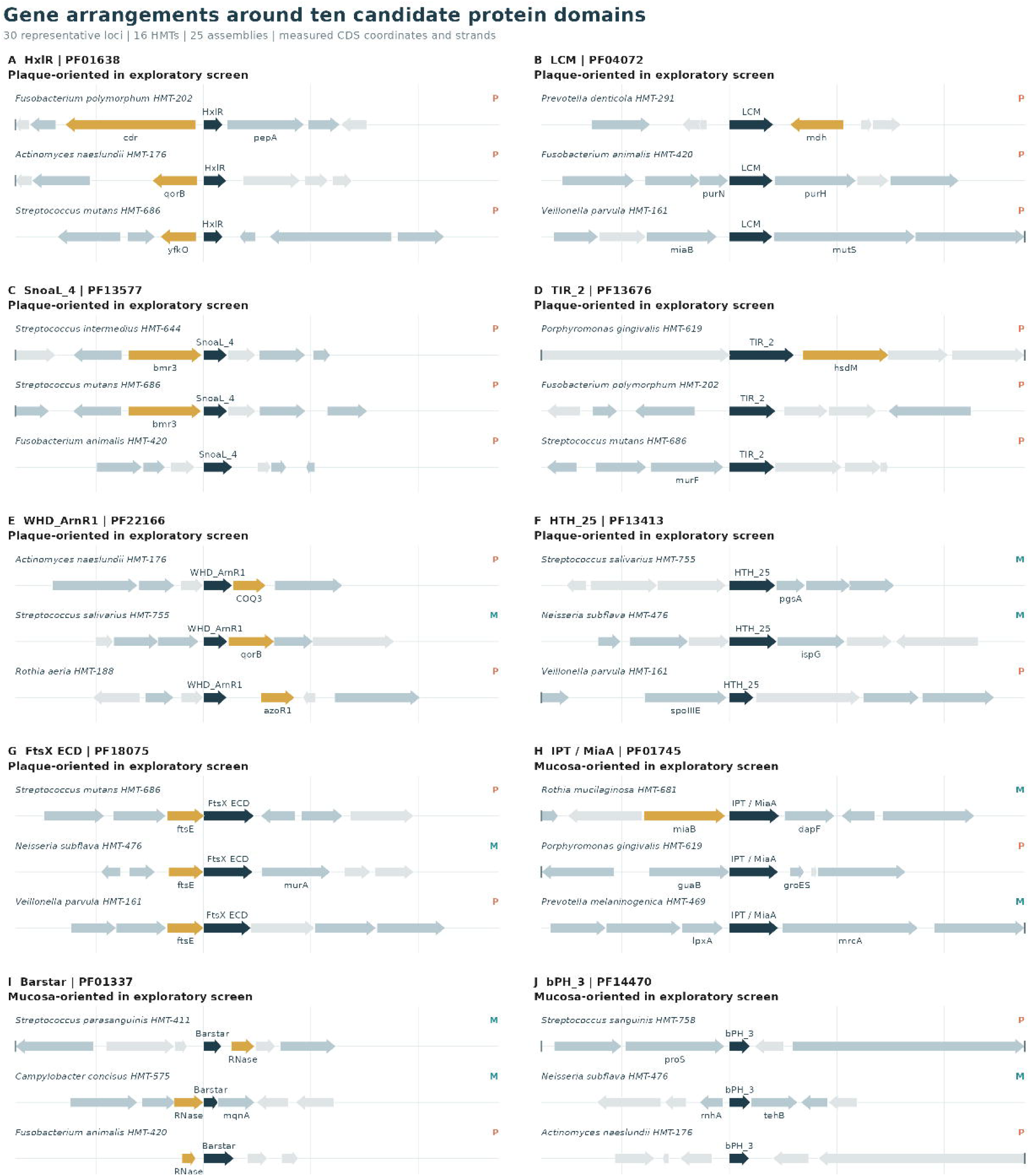
Sister-pair genomic contrasts provide a local phylogenetic sensitivity analysis. Eight terminal sister pairs exhibiting opposite plaque–mucosa affinities were identified in the 47-HMT marker phylogeny. For each genus, points show plaque-minus-mucosa prevalence differences for PF04072 (LCM; circles) and PF12680 (SnoaL_2; diamonds). Node-support values are reported for the terminal relationships; the *Gemella* pair received 59% bootstrap support and the remaining seven 100%. LCM favoured the plaque-associated sister in six pairs, favoured the mucosal sister in *Porphyromonas* and was tied in one; SnoaL_2 favoured plaque in six pairs and was tied in two.

## Data availability

Reference genomes and ecological abundance summaries were obtained from the public resources described in Methods. The accompanying data package contains the accession-level manifest, HMT-to-assembly assignments, ecological scores, nonredundant feature matrix and figure-source tables.

## Code availability

The accompanying code package contains the analysis scripts, workflow templates, software requirements and R figure-generation scripts: https://github.com/AbishStephen/oral-microbial-habitat-affinity.

## Acknowledgements

This research utilised Queen Mary’s Apocrita HPC facility, supported by QMUL Research-IT. http://doi.org/10.5281/zenodo.438045

## Ethics approval

This study reanalysed publicly available microbial genome and environmental abundance datasets and did not recruit new human participants or collect identifiable participant data.

